# Disrupted object-scene semantics boost scene recall but diminish object recall in drawings from memory

**DOI:** 10.1101/2020.05.12.090910

**Authors:** Wilma A. Bainbridge, Wan Y. Kwok, Chris I. Baker

**Affiliations:** Department of Psychology, University of Chicago, Chicago, IL 60637; Laboratory of Brain and Cognition, National Institute of Mental Health, Bethesda, MD 20814

**Author notes:** indicates equal contribution (Wilma A. Bainbridge, Wan Y. Kwok). <u>Corresponding author:</u> Wilma A. Bainbridge, Department of Psychology, University of Chicago, 5848 South University Ave, 303 Beecher Hall, Chicago, IL 60637.

**Keywords:** Saliency, binding errors, global scene processing, local scene processing

## Abstract

Humans are highly sensitive to the statistical relationships between features and objects within visual scenes. Inconsistent objects within scenes (e.g., a mailbox in a bedroom) instantly jump out to us, and are known to capture our attention. However, it is debated whether such semantic inconsistencies result in boosted memory for the scene, impaired memory, or have no influence on memory. Here, we examined the relationship of scene-object consistencies on memory representations measured through drawings made during recall. Participants (N=30) were eye-tracked while studying 12 real-world scene images with an added object that was either semantically consistent or inconsistent. After a 6-minute distractor task, they drew the scenes from memory while pen movements were tracked electronically. Online scorers (N=1,725) rated each drawing for diagnosticity, object detail, spatial detail, and memory errors. Inconsistent scenes were recalled more frequently, but contained less object detail. Further, inconsistent objects elicited more errors reflecting looser memory binding (e.g., migration across images). These results point to a dual effect in memory of boosted global (scene) but diminished local (object) information. Finally, we replicated prior effects showing that inconsistent objects captured eye fixations, but found that fixations during study were not correlated with recall performance, time, or drawing order. In sum, these results show a nuanced effect of scene inconsistencies on memory detail during recall.

## Introduction

When we view a scene, we automatically parse many aspects of that scene—its overall gist, constituent objects, and their relations to each other and the greater scene layout (Oliva and Torralba, 2006; Fei-Fei, Iyer, Koch, & Perona, 2007). We are highly sensitive to the meanings of objects within scenes (Henderson & Hayes, 2017), and form expectations on how objects relate to their scene backgrounds with what can be considered a “scene grammar” (Võ, Boettcher, & Draschkow, 2019). Thus, when we see scenes containing violations of this grammar—for example, when a beach ball is unexpectedly in a laboratory—we immediately notice. Such inconsistencies in object-scene semantics cause disruptions in our ability to process these images (Greene, Botros, Beck, & Fei-Fei, 2015), and have been shown to capture eye movements during perceptual and visual search tasks (Loftus & Mackworth, 1978; De Graef, Chrisstiaens, & d’Ydewalle, 1990; Henderson, Weeks, & Hollingworth, 1999; Malcolm & Henderson, 2010), even when performing an irrelevant task (Cornelissen & Võ, 2017).

However, even though observers fixate longer on inconsistent objects, it is unclear how scene consistency influences the later memory for that scene and its objects. While some research has reported no memory differences between inconsistent and consistent objects in scenes (Cornelissen & Võ, 2017), other work has reported boosted memory for inconsistent objects (Friedman, 1979; Pedzek, Whetstone, Reynolds, Askari, & Dougherty, 1989; Hollingworth, Williams & Henderson, 2001) or, conversely, boosted memory for consistent objects (Draschkow & Võ, 2017). Given these divergent results, how do scene semantics shape the memory representations for a scene? Further, what specific aspects of a scene’s memory are impacted by its semantics – memory for the entire image, the manipulated object, or the background scene?

In the current study, we compare the underlying visual memory representations for inconsistent and consistent scenes using a visual recall drawing task. Previous work assessing the influences of scene semantics on memory have relied on verbal recall or visual recognition tasks (Friedman, 1979; Pezedk et al., 1989; Hollingworth et al., 1998; Cornelissen & Võ, 2017). However, these types of measures may provide limited information about the nature of a memory – only revealing whether an item is remembered or not, but not what specific visual content of that memory drives its recollection. Recent work has discovered that using drawing-based recall to quantify memory can reveal more fine-grained information than verbal recollection, and requires no assumptions about matched foil images, often necessary for visual recognition tasks (Bainbridge, Hall, and Baker, 2019). Thus, such a drawing task may reveal subtler differences between memories for consistent and inconsistent scenes than was possible to capture in previous work. Further, drawings can be objectively quantified through online crowd-sourced scoring to reveal a wide range of information, including object detail, spatial accuracy, and inclusion of false additional objects (Bainbridge et al., 2019). With such a task, we can thus examine not only whether an inconsistent object is remembered better than a consistent object, but how inconsistency impacts memory for other objects in the scenes, their spatial relations, and the scene overall.

With these measures, we examined several questions for how scene semantics might influence memory representations. First, we consider how consistency affects recall of the manipulated object in the scene. One possibility is that inconsistent objects are distinctive and easier to remember (Friedman, 1979; Pedzek, Whetstone, Reynolds, Askari, & Dougherty, 1989; Hollingworth, Williams & Henderson, 2001). Conversely, we might instead find that consistent objects better fit our scene schemas and thus are easier to remember, as has been found in recent work analyzing the role of consistency on scene construction (Draschkow & Võ, 2017). A third possibility is that we may observe no memory difference between inconsistent and consistent objects (Cornelissen & Võ, 2017). Second, we consider how consistency affects memory for the overarching image – do participants tend to draw inconsistent or consistent images more frequently from memory, regardless of their memory for the manipulated object? Finally, beyond memory for the inconsistent / consistent object or its encompassing image, we ask whether there is a difference in memory for the other objects in the scene (what we will hereby refer to as the “background scene”). On one hand, the heightened distinctiveness of an image owing to the presence of an inconsistent object could boost memory for the entire scene, including surrounding objects. On the other hand, these inconsistent objects could create a “spotlight” effect, capturing attention away from surrounding objects (Cornelissen & Võ, 2017) and reducing recognition for objects semantically unrelated to the inconsistent object (Auckland, Cave, & Donnelly, 2007; Davenport, 2007). With this spotlight effect, we might also observe transpositions of objects that are semantically unrelated to their overarching image (Hannigan & Reinitz, 2003). Thus, by analyzing drawings made from memory, we can examine memory performance for the image, the manipulated object, as well as the background scene. Further, using both eye-tracking and computer-vision based saliency models during image encoding as well as pen-tracking during drawing recall, we can see whether we can replicate previous findings on increased fixations to inconsistent objects (e.g., Cornelissen & Võ, 2017), and whether fixation patterns during perception predict recall performance.

To preview our results, we find an interesting trade-off in memory, in which semantically inconsistent images are recalled more frequently, but with less detail, and with weaker binding between the inconsistent object and the scene, resulting in transpositions of that object across images. Further, while we replicate the observation that inconsistent objects capture attention during encoding, we find that recall patterns cannot be explained by fixation patterns or image saliency during encoding.

## Methods

### Participants

Thirty adults (9 male, 21 female) were recruited from the local Washington D.C. area for participation in this experiment. This sample size was derived from a previous, similar drawing-based experiment (Bainbridge et al., 2019). Participants were healthy native English speakers with corrected or normal vision. All participants consented following the guidelines of the National Institutes of Health (NIH) Institutional Review Board (NCT00001360, 93M-0170) and were compensated for their participation. 1,725 online scorers were recruited from online crowd-sourcing task platform Amazon Mechanical Turk (AMT), acknowledging their participation following the guidelines of the NIH Office of Human Subjects Research Protections (OHSRP), and were also compensated for their participation.

### Stimuli

Stimulus images were created from twelve distinctive scene images from different scene categories, half indoor (bathroom, bedroom, classroom, kitchen, laboratory, laundry room) and half outdoor (campsite, construction site, neighborhood street, playground, swimming pool, backyard). Adobe Photoshop was used to naturally add an object to each image (referred to throughout as the “manipulated object”) that was either consistent or inconsistent with the scene semantics (Figure 1). The scene images were paired, and these object manipulations were conducted within the pairs, so that the consistent object in a given image was also used as the inconsistent object in its paired image, and vice versa. For example, in the consistent condition, a lab scene contained a microscope and a pool scene contained a beach ball (Figure 1). In the inconsistent condition, the lab scene had a beach ball and the pool scene had a microscope. This resulted in a set of 24 stimuli, comprising of a consistent and inconsistent version of each of the twelve scenes (and, similarly, each of twelve objects had a consistent image and an inconsistent image). To confirm that we successfully manipulated scene consistency, all images were rated online by Amazon Mechanical Turk (AMT) workers (N=15 per image; N=67 total) on a 5-point Likert Scale on how typical (“normal”) it was for that object to be in the scene (1 = Very Abnormal, 5 = Very Normal). As expected, consistent objects were rated to be significantly more normal than inconsistent objects (Consistent: M=4.4 SD=1.1; Inconsistent: M=1.6 SD=1.1; Wilcoxon signed rank test: *Z*=2.20, *p*=0.028). During the main experiment, each participant saw 12 images (one of each background scene), with half consistent images and half inconsistent images. Which images were consistent or inconsistent was pseudo-randomly counterbalanced across participants, so each of the 24 images was ultimately seen by fifteen participants.

**Figure 1 –.**
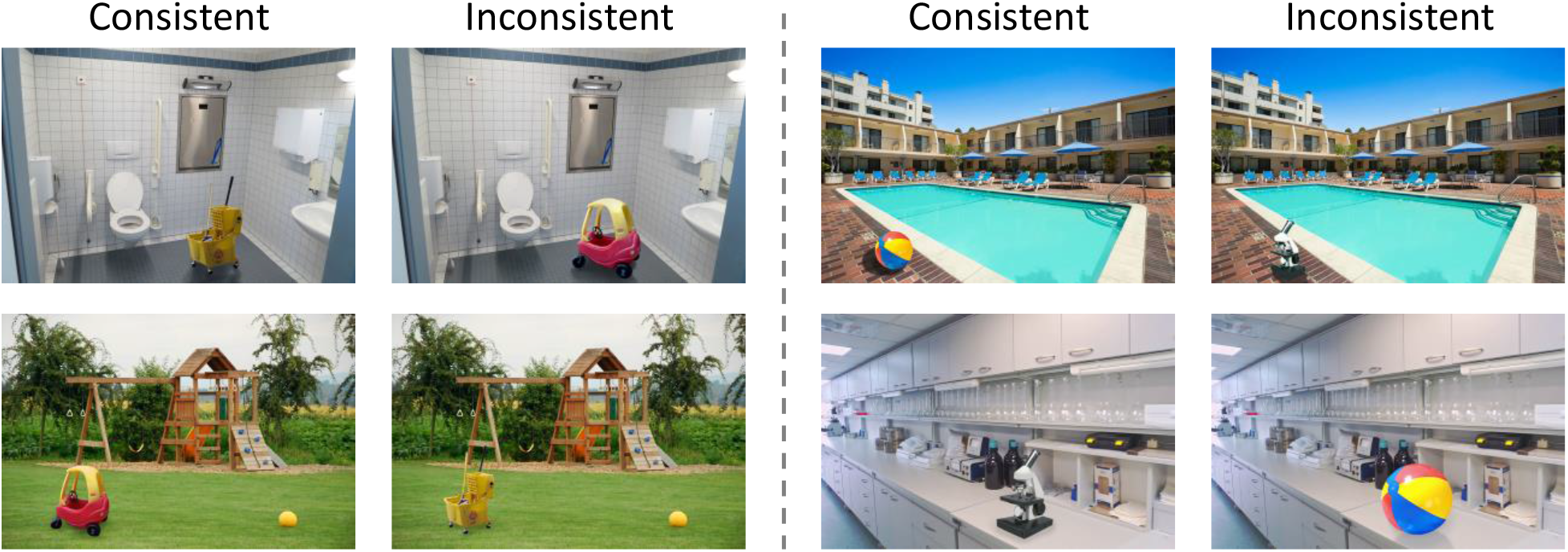
Two example sets of consistent and inconsistent scenes. (Left) A toy car or mop bucket in a bathroom or playground. (Right) A beach ball or a microscope on a swimming pool deck or in a laboratory.

### Experimental Procedures

At the beginning of the experiment, participants were told to carefully examine each image as they would be later tested on their memory. However, participants were unaware of the nature of the memory task, and were unlikely to expect a drawing task. The experiment was split into four phases (Figure 2).

**Figure 2 –.**
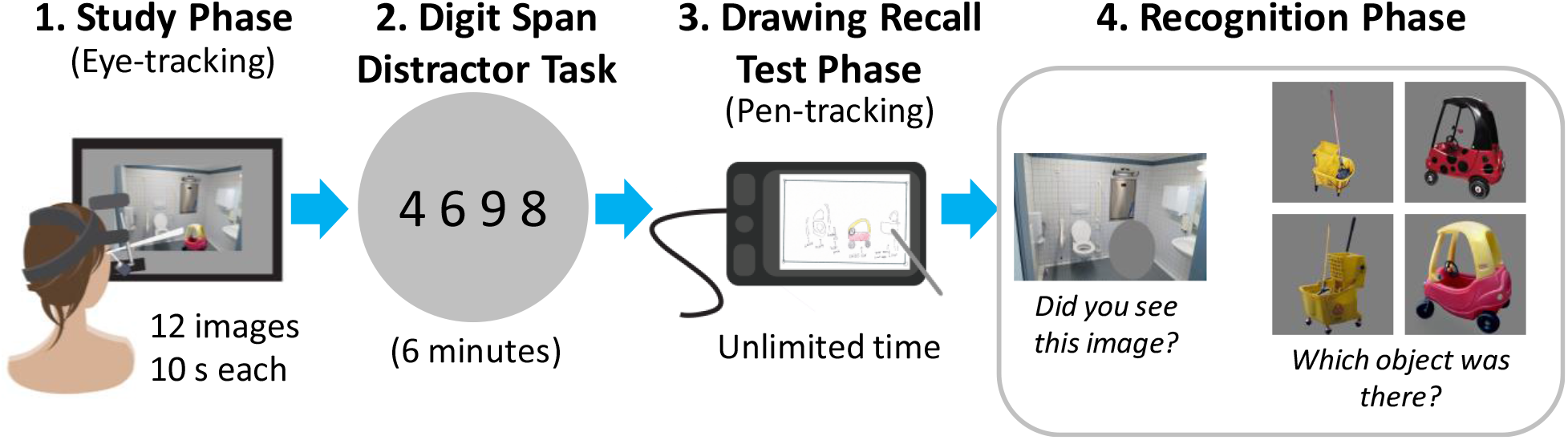
The experimental procedures. The experiment consisted of four phases: 1) A study phase in which participants studied 12 randomly ordered images (6 consistent, 6 inconsistent) for 10s each while their fixations were tracked, 2) A digit span distractor task in which participants had to verbally recall digit series, 3) A drawing recall test phase in which participants drew the studied scenes from memory while their pen movements were tracked, and 4) A recognition phase in which participants had to separately recognize the scene and the manipulated object.

**Figure 3 –.**
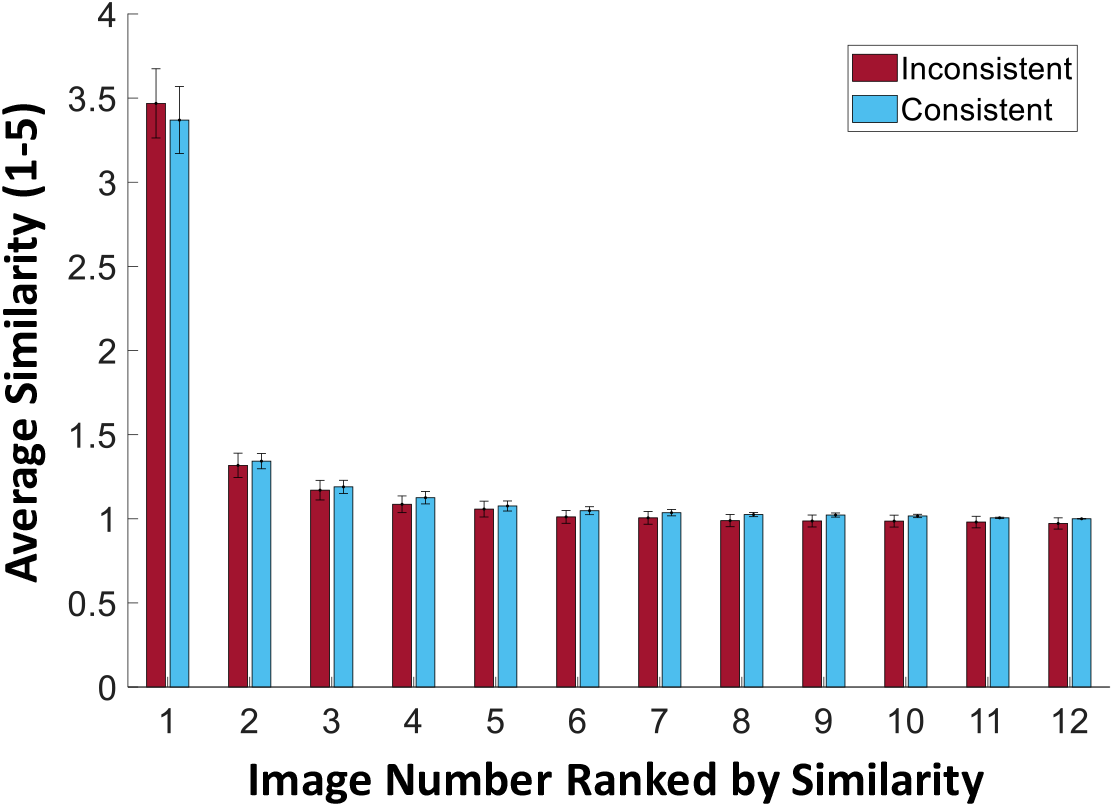
Diagnosticity of the drawings across images for the two conditions. The average ratings made by online scorers of the similarity of participants’ drawings to each of the 12 images they saw, ranked from highest to lowest. Similarity here was assessed on a scale of 1 (low) to 5 (high) with a question asking how likely it was that a drawing was of a given image. Red bars indicate drawings made of inconsistent images, while blue bars indicate drawings made of consistent images. The high spike for image #1 and quick drop-off for images #2-12 (all averaging below a rating of 1.5) for both conditions indicates that it was clear to AMT scorers that a given drawing was highly similar to only one image and dissimilar from all others. In other words, drawings were highly diagnostic of their images. There was no significant difference in diagnosticity between consistent and inconsistent images. Error bars indicate standard error of th emean.

The first phase was a study phase, in which participants viewed each of the twelve images for 10 seconds while their eye movements were tracked with a head-mounted EyeLink 1000 Plus eye-tracking device. Between the presentation of each stimulus image, a fixation cross was displayed to the right of the image on the screen, in order to avoid biasing eye movements to the center. After the participant fixated on the cross, the next stimulus was then displayed. Each image was displayed at 1200 x 800 pixels on a 1920 × 2000 resolution 24-inch screen.

The second phase was a digit span distractor task intended to disrupt verbal working memory strategies. Participants saw a consecutive series of digits varying by 3-9 digits in length, and then had to repeat back the series of digits from memory when prompted. This repeated for 21 trials, and introduced an approximately 6-minute delay between the study and test phases.

The third phase was the drawing recall test phase. Participants were given sheets of paper with a rectangular outline with dimensions matching those of the original images, and were asked to draw as many images as they could remember in as much detail as possible. Participants drew on a Wacom Paper Pro tablet, which allowed participants to draw with an inked pen on paper while it simultaneously recorded pen strokes digitally in real time. Participants were told to draw the images in any order. They were also given colored pencils if they wanted to include color detail in their drawings, but were asked to include color only if they specifically recalled it. They were instructed to add color after completing the pen drawing of all the objects they recalled. Participants were also told they could label objects if they wished to clarify what they were. Participants were given as much time as they needed and took 27 minutes on average for the recall phase (SD=8).

In the fourth and final phase, participants completed a recognition phase, in which they made a series of recognition-based judgments of the images. They were shown the 12 scene images they studied randomly interspersed with 12 closely-matched foil scenes of the same scene categories. All scene images had a gray occluding ellipse covering the manipulated (consistent or inconsistent) object. Foil images had a gray occluding ellipse placed in a plausible similar location. First, for a given image, participants were asked if they had seen the scene during the study phase (scene recognition). If they said yes, they were presented with four object images and had to indicate which was the object they saw in that scene. The four choices of object images were: 1) the inconsistent object, 2) the consistent object, 3) a different exemplar from the inconsistent object category, and 4) a different exemplar from the consistent object category. This question tested both object category recognition (e.g., if you studied an inconsistent scene, did you falsely remember seeing a consistent object?), as well as specificity to the exemplar within the same category (e.g., did you remember that you saw that specific microscope in the scene, or a microscope in general?).

### Online Scoring Procedures

The resulting 275 drawings were scanned and uploaded to AMT, to crowd-source worker ratings on several properties of the memory drawings. Specifically, four different sets of measures were collected for each drawing. For all rating tasks, each AMT worker could participate in as many trials as they wanted.

#### Drawing Match Scoring

AMT workers rated how well each drawing matched one of the images participants studied, giving us a measure of diagnosticity of that drawing. For a given trial, they were presented with a drawing with an image next to it and rated on a 5-point Likert scale the likelihood that it was a drawing of that image (1= Definitely Not, 5 = Definitely). Across trials, drawings were tested against each of the 12 images seen by that participant. Twelve ratings were collected for each drawing, and 611 AMT workers participated in total. The majority response was selected as the corresponding image (“original image”) for that drawing.

#### Object Identification Scoring

AMT workers determined which objects were included in each drawing. For a given trial, they were presented with a drawing and five copies of the matched original image with a different object highlighted on it. AMT workers had to click on the objects from the image that were present in the drawing. Five AMT workers rated each object and 679 AMT workers participated in total. Objects were determined to be in the drawing if at least three out of five workers said it was in the drawing.

#### Object Location Scoring

AMT workers determined the locations and sizes of each object present in the drawings. For a given trial, they were presented with a drawing and the original image with an object highlighted on it. On the drawing, they had to place and resize an ellipse to encircle that specified object. Five AMT workers made ellipses for each object and 453 AMT workers participated in total. The final ellipse was determined by the median centroid and radii in the x and y directions.

#### False Additional Object Scoring

AMT workers determined the presence of additional objects in the drawings that were not in the original images. For a given trial, they were presented with a drawing and its corresponding image and had to write down all objects that existed in the drawing but not the image. Fifteen AMT workers rated each image and 200 AMT workers participated in total. Any objects listed by at least five workers were counted as false alarms.

### Fixation, Pen-tracking, and Saliency Analyses

Before running the experiment, we segmented the outlines of all objects within each of the 24 images using the online object annotation tool LabelMe (Russell, Torralba, Murphy, & Freeman, 2008). From the EyeLink 1000 Plus, we extracted fixation-based heatmaps for each participant to each image. To obtain a metric of fixation time per object, we computed a normalized fixation time, determined by the total fixation time across all pixels within a given object, divided by the total fixation time across all pixels within the image. We also looked at fixation order by comparing the order in which the manipulated object had its first fixation in relation to the first fixation on all other objects (e.g., was the manipulated object fixated before other objects?). Fixation order was then normalized by total number of objects in the drawing.

For recall, we calculated object-based recall scores as the proportion of participants who drew each object, divided by the number of participants who drew the image containing that object. Using the tablet recordings of the pen movements, we also calculated amount of time spent drawing each manipulated object. A scorer watched the video of pen strokes created by the drawing tablet for each drawing. The start and end time of pen strokes for the manipulated objects were noted for each image. Total amount of time spent drawing the object was calculated as the difference between the end time and the start time, normalized by total amount of time spent on the drawing. Similarly, we calculated sequential drawing order by assigning an order to each object based on first pen stroke on that object. Drawing order was then normalized by total number of objects in the drawing. One participant was removed from the drawing time and drawing order analyses due to a technical glitch with the pen tablet software (resulting in N=29 for these analyses).

To compute image saliency scores, we used two state-of-the-art computer vision algorithms designed to predict human fixation patterns: DeepGaze II (Kümmerer, Wallis, & Bethge, 2016) and Graph-Based Visual Saliency (Harel, Koch, & Perona, 2007). DeepGaze II is a model that predicts fixation patterns based on features from the VGG-19 deep neural network for object identification combined with a readout network trained for saliency prediction based on human fixations (Kümmerer et al., 2016). Graph-Based Visual Saliency (GBVS) is a model that identifies visually dissimilar regions of an image (Harel et al., 2007). For both metrics, we obtained saliency heatmaps for each of the stimulus images (see Figure 7). Object-based saliency was then calculated as the average saliency across the pixels of that object, normalized by the average saliency of the entire image.

### Data Analyses

For most analyses, we conducted paired samples t-tests within subjects to compare the above metrics between participants’ drawings of consistent scenes versus inconsistent scenes. We first tested these metrics for normality using a Kolmogorov-Smirnov goodness-of-fit test, and found none of these were significantly different from a normal distribution (all *p*>0.05). For metrics with limited ranges (e.g., Likert scales, number of drawings made), we instead conducted non-parametric paired samples Wilcoxon signed rank tests.

## Results

### Drawings are highly diagnostic of their images

The first step is identifying what image each drawing corresponds to. Further, given the range of people’s drawing abilities and memory, can a separate group of participants tell what image a drawing represents? AMT workers saw individual drawings matched with each of the 12 images studied by participants in the main experiment, and judged the likelihood that the drawing was of that image on a scale of 1 (Definitely Not) to 5 (Definitely). Overall, it was clear to AMT workers what images matched the drawings, with only a single image getting a score above 3 on average (Figure 2). For all further analyses, the highest rated image was taken as the corresponding image for each drawing. Importantly, there was no significant difference in ratings between consistent and inconsistent images (Consistent: M=3.9, SD=0.5; Inconsistent: M=3.7, SD=0.6; Wilcoxon signed rank test: *Z*=1.70, *p*=0.090). This means that both semantically consistent and inconsistent drawings could be matched with their original images, and were equally diagnostic of their original image. However, this rating of diagnosticity serves as a relatively coarse metric, as several different features could contribute to being able to successfully match a drawing to an image (i.e., the manipulated object, properties of the background scene). A closer look at the content in these drawings may reveal key differences between consistent and inconsistent images.

### More inconsistent scenes are recalled than consistent scenes

The memory drawing experiment resulted in 275 total drawings, with 126 drawings of consistent images, and 149 drawings of inconsistent images (Figure 4). This reflects a general tendency to recall inconsistent images over consistent images (Chi-squared test for proportions: *χ*^2^=3.85, *p*=0.050). Each participant on average drew 9.2 drawings from memory out of the 12 that they studied (SD=2.16, Min=5, Max=12). Of those drawings, participants drew more inconsistent images than consistent images from memory (Wilcoxon signed rank test: *Z*=2.10, *p*=0.036), drawing on average 5.0 inconsistent images (SD=1.5) and 4.2 consistent images (SD=1.5). Thus, memory for inconsistent images overall was better than that for consistent images.

**Figure 4 –.**
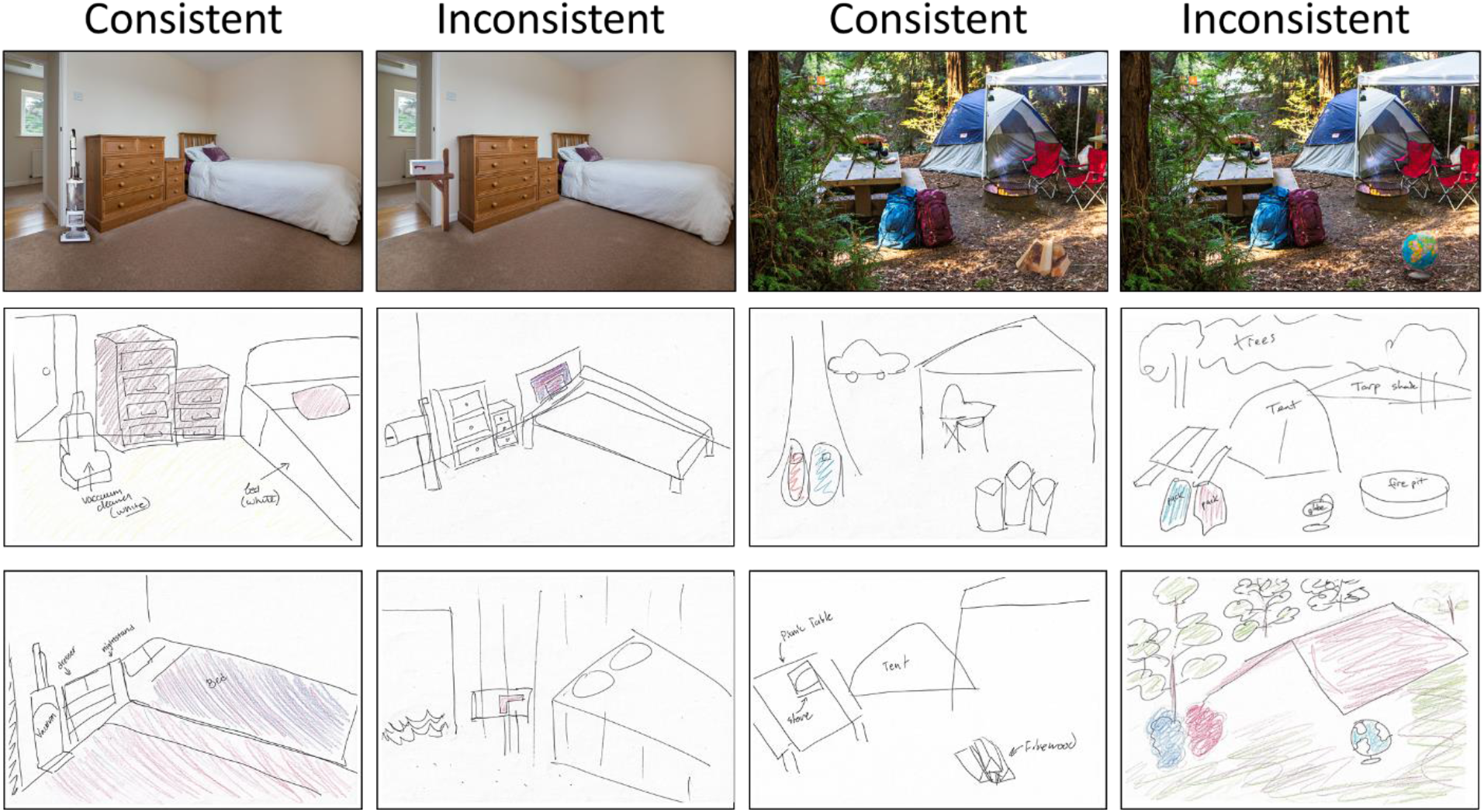
Example drawings for four of the stimulus images. Two example drawings each for the consistent and inconsistent bedroom scene, and the consistent and inconsistent camp scene. Each drawing was taken from a different participant, showcasing the impressive level of both object and spatial detail in the memory drawings for a range of people. The key question in this study is whether there are differences in memory detail between drawings for the consistent and the consistent scenes.

### More objects are recalled in consistent scenes than inconsistent scenes

Next, we looked at the amount of detail available in each drawing by having AMT workers judge which objects from the original image were included in each drawing. Each image contained on average 39.2 objects (with the same number of objects across the consistent and inconsistent versions of a given scene), and participants drew on average 9.0 objects per image (SD=2.9), or 77.6 objects on average across the experiment. Participants drew a significantly higher proportion of the objects in consistent drawings versus inconsistent drawings (Consistent: M=23.4%, SD=6.9%; Inconsistent: M=19.8%, SD=8.1%; *t*(28)=2.44, *p*=0.021), as well as if the manipulated object is excluded (*t*(28)=2.56, *p*=0.016). We then looked to see whether there were differences in the tendency to draw the manipulated object (the consistent or inconsistent object). We found no significant difference between consistent or inconsistent drawings in the proportion containing the manipulated object (Consistent: M=53.8%, SD=25.8%; Inconsistent: M=65.0%, SD=28.1%; *t*(28)=1.37, *p*=0.181). This indicates that while semantically inconsistent objects were recalled just as frequently as their consistent counterparts, there was reduced memory for objects in the inconsistent background scenes than consistent ones.

### The nature of object errors in consistent versus inconsistent scenes

We then investigated whether there were differences between conditions in the types of object errors that were made in the drawings. Overall, participants included relatively few false additional objects in their drawings, only drawing 25 objects that didn’t exist in the consistent images (across 126 drawings), and 21 objects that didn’t exist in the inconsistent images (across 149 drawings). Within participants, there was no significant difference in the number of false additional objects they drew for inconsistent scenes versus consistent scenes (*t*(29)=0.64, *p*=0.526). Thus, differences in scene semantics did not appear to induce differences in false memories in these drawings.

However, participants made intriguing errors with the manipulated object in their drawings (see Figure 5). In 18 drawings (13 inconsistent, 5 consistent), participants made drawings of only the manipulated object, unable to recall the surrounding background scene. In 6 drawings (6 inconsistent, 0 consistent), participants drew a detailed scene and included a circle with an unspecified object; they remembered that the manipulated object was there, but not what it was. Finally, in 16 drawings (13 inconsistent, 3 consistent), participants transposed the manipulated object so that it was in a different scene they had viewed. All of these errors occurred significantly more frequently for inconsistent than consistent scenes (Chi-squared test of proportions, Isolated Object: *χ*^2^=7.11, *p*=0.008; Unspecified Object: *χ*^2^=12.00, *p*=5.32 × 10^-4^; Transposed Object: *χ*^2^=12.50, *p*=4.07 × 10^-4^). These results imply that a disruption of scene semantics may result in a looser binding in memory of that inconsistent object with its encompassing scene.

**Figure 5 –.**
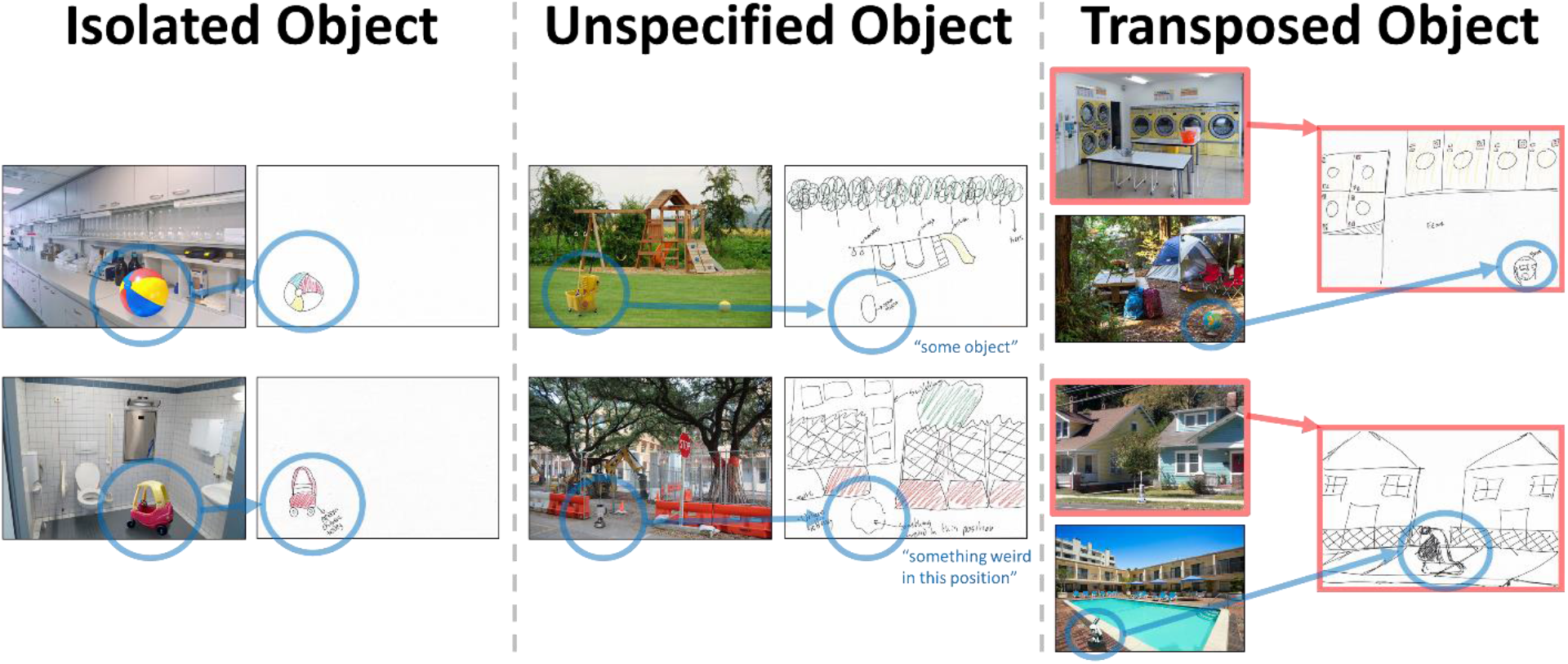
Examples of memory errors made by participants for the manipulated object. Each example is taken from a different participant. We identified three types of errors: 1) drawing the object in isolation (left), 2) drawing a detailed scene with a circle noting recollection of an unspecified object at that location but not its identity, and 3) transposing the object to a different scene. For the unspecified object errors, the text labeling the circle is included in larger font. These errors occurred overwhelmingly more often when objects were inconsistent with their background scenes (*p* < 0.01 for all error types).

### Equally high spatial accuracy in consistent and inconsistent images

While there are differences in object memory based on scene semantics, are there also differences in spatial accuracy for the objects in the drawings? AMT workers indicated the size and location of each object in the drawing by drawing an ellipse on the drawing. With that ellipse, we calculated mean location error (centroid x and y error) and size error (radius x and y error) for each object. In both conditions, low amounts of spatial error were found, although errors were larger in magnitude in the Y-direction than the X-direction. Errors of object location were transpositions of less than 11% of the size of the entire image (X-direction: Consistent M=2.2%, Inconsistent M=0.4%, Y-direction: Consistent M=9.3%, Inconsistent M=10.8%). Errors in size were on average less than 4% image’s pixels (Width: Consistent M=2.2%, Inconsistent M=2.0%; Height: Consistent M=2.9%, Inconsistent M=3.7%). There was no significant difference between consistent and inconsistent drawings in spatial accuracy, neither in terms of location accuracy (X-direction: *t*(28)=1.45, *p*=0.157; Y-direction: *t*(28)=0.66, *p*=0.512), nor object size (Width: *t*(28)=0.35, *p*=0.732; Height: *t*(28)=1.43, *p*=0.163). These results indicate that manipulations of object semantics do not appear to affect spatial accuracy in memory.

### Comparing eye fixation patterns, visual saliency, and recall

Prior studies have observed an influence of scene semantics on fixation patterns (Loftus & Mackworth, 1978; De Graef, Chrisstiaens, & d’Ydewalle, 1990; Henderson, Weeks, & Hollingworth, 1999; Malcolm and Henderson, 2010). We examined whether the inconsistent objects in our paradigm captured attention. Further, we investigated whether these fixation patterns (Figure 6) could be predicted by visual saliency of the image. Finally, we tested the degree to which these metrics related to recall of the information in the images.

**Figure 6 –.**
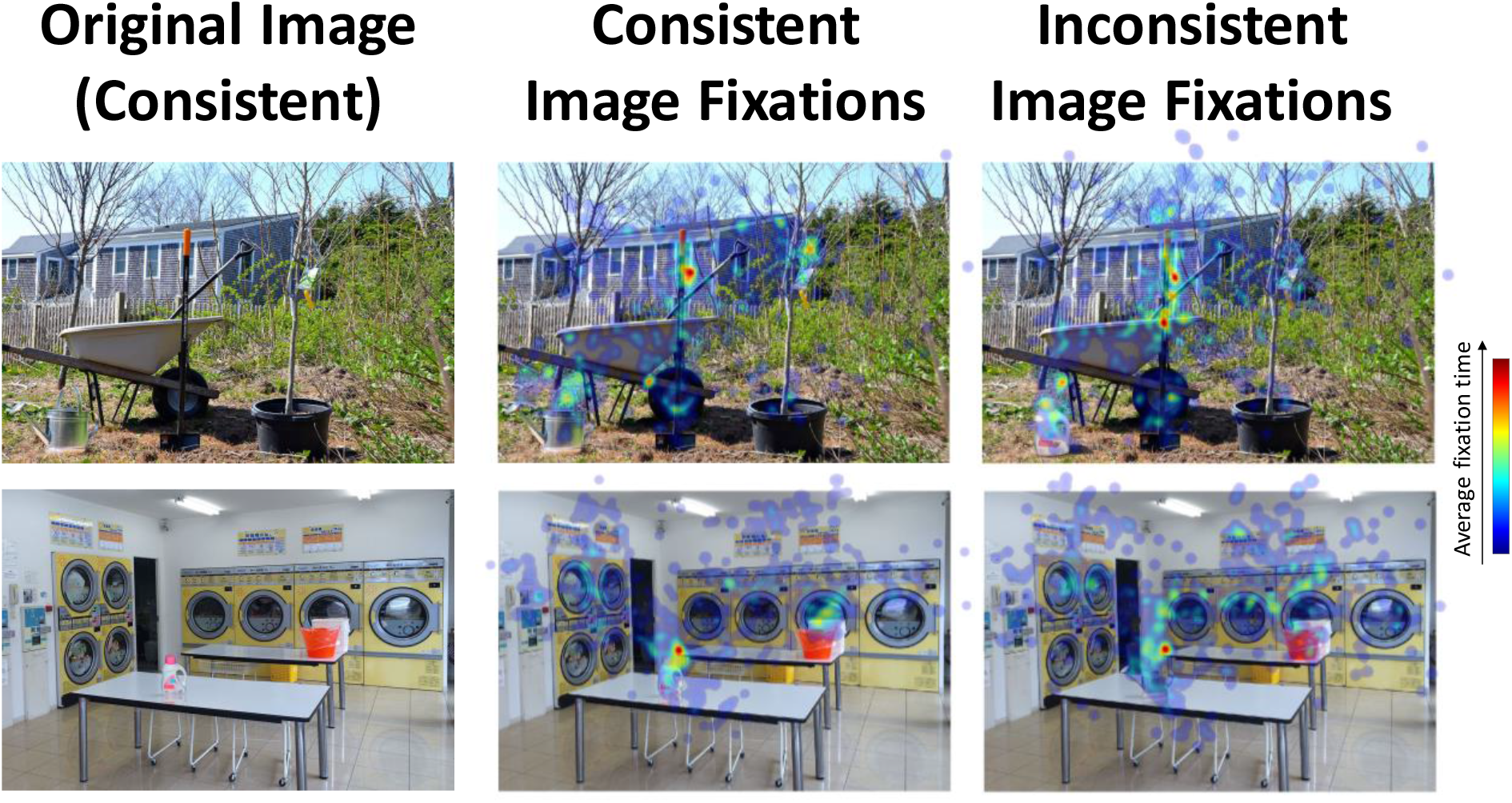
Example average fixation heatmaps. Heatmaps showing examples of average fixation patterns (averaged across participants) for the consistent and inconsistent versions of two paired scenes where their objects were swapped (i.e., a watering can or laundry detergent in a backyard scene or a laundry scene). Red indicates higher total fixation time on average, and blue indicates lower total fixation time. For the backyard scene, the inconsistent detergent bottle causes more fixations than the consistent watering can. For the laundry scene, both the watering can and detergent bottle elicit more fixations.

We looked at fixation time for each object during study, normalized by total amount of time spent fixating on the image. Participants fixated for significantly longer on the inconsistent manipulated object than the consistent manipulated object (Inconsistent: M=11.29% of fixation time, SD=7.27%; Consistent: 6.91% of fixation time, SD=6.34%; *t*(29)=3.51, *p*=0.002). We then looked to see whether current state-of-the-art visual saliency algorithms DeepGaze II and GBVS could predict these fixation patterns (see Methods). We found a significant correlation between DeepGaze-predicted saliency and fixation time on the manipulated objects (Spearman’s rank correlation: *ρ*=0.634, *p*=0.001) as well as the same correlation for GBVS-predicted saliency and fixation time (*ρ*=0.634, *p*=0.001). These results indicate that visual saliency may be able to partially account for greater fixations to inconsistent objects; we discuss these implications later in the Discussion.

Next, we investigated whether these metrics could predict the proportion of people who recalled the manipulated object (Figure 7). We observed no significant correlation between mean fixation time and recall proportion (*ρ*=0.032, *p*=0.881). We also observed no significant correlation between recall proportion and DeepGaze-predicted saliency (*ρ*=0.137, *p*=0.524), nor GBVS-predicted saliency (*ρ*=0.201, *p*=0.346). Thus, while inconsistent objects may capture eye-movements and be more visually salient, neither fixation patterns nor image-based saliency can account for differences in memory performance between the consistent and inconsistent objects.

**Figure 7 –.**
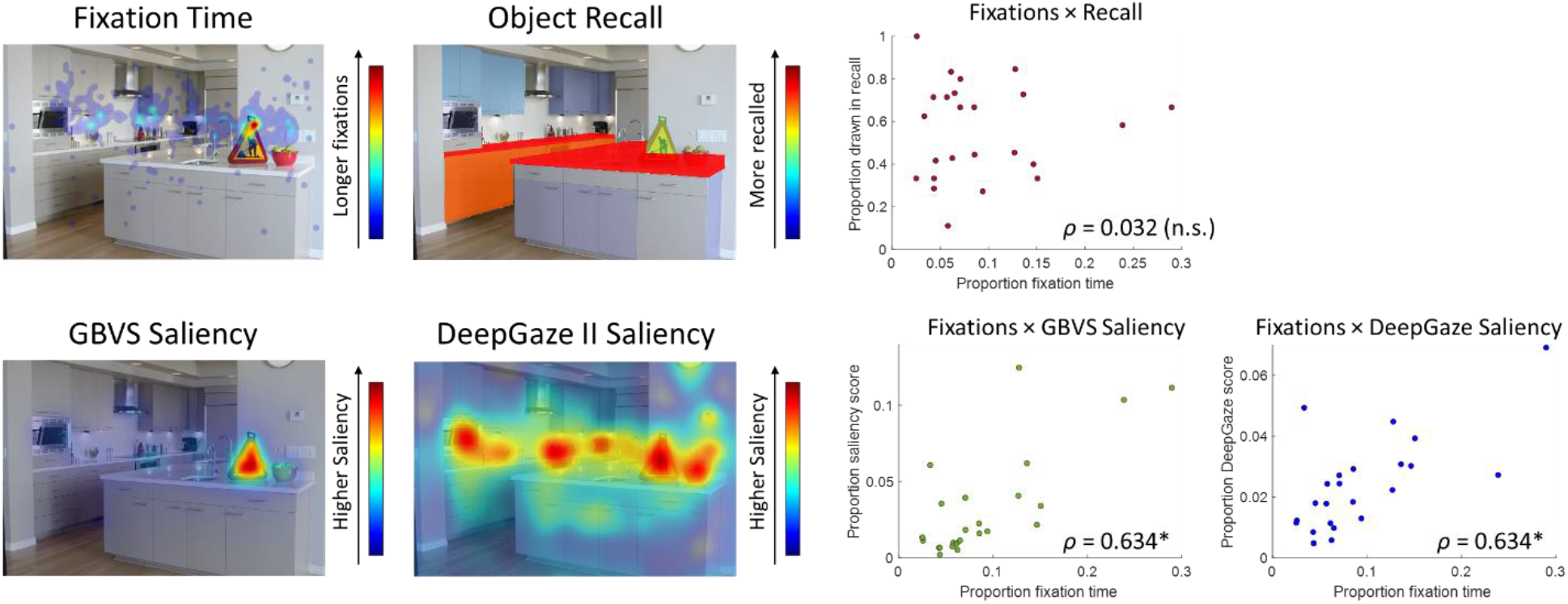
Comparison of fixation time during study, visual saliency, and recall success. (Left) For each stimulus image, we generated four different heatmaps: 1) eye fixation times across the pixels of the image, 2) proportion of participants recalling each object in the image, 3) visual saliency of the image calculated using Graph-Based Visual Saliency (Harel et al., 2007), and 4) visual saliency of the image calculated using DeepGaze II (Kümmerer et al., 2016). (Right) Scatterplots of average fixation time with the three other metrics (recall proportion, GBVS visual saliency, and DeepGaze II visual saliency). Each point represents one of the 24 stimulus images, and indicates the average score for the manipulated object. While saliency metrics were significantly correlated with fixation time, no measure was significantly correlated with recall success. Correlations reported here are Spearman’s *ρ,* with * indicating significant correlations.

### Comparing the temporal order of recall for consistent and inconsistent scenes

In conjunction with recording eye movement patterns during study of the images, we also recorded real-time pen movement patterns during recall of the images (Figure 8). Participants did not spend a significantly different amount of time drawing the inconsistent versus consistent manipulated objects (Inconsistent: M=16.08 s, SD=7.75; Consistent: M=15.85 s, SD=11.26; *t*(27)=0.07, *p*=0.942), nor a significantly different amount of time drawing inconsistent versus consistent images (Inconsistent: M=2.03 min, SD=0.62; Consistent: M=2.28 min, SD=0.89; *t*(27)=1.53, *p*=0.139). There was also no significant difference in the order in which inconsistent versus consistent objects were drawn (*t*(25)=1.01, *p*=0.323). It was thus not the case that inconsistent objects were drawn for longer or drawn earlier. There was also no significant correlation between amount of time spent fixating the manipulated object and amount of time drawing the manipulated object (Spearman’s rank correlation: *ρ*=0.205, *p*=0.130). Thus, time spent during the drawing recall phase does not reveal clear differences between the inconsistent and consistent images, nor a clear relationship to fixation patterns during study.

**Figure 8 –.**
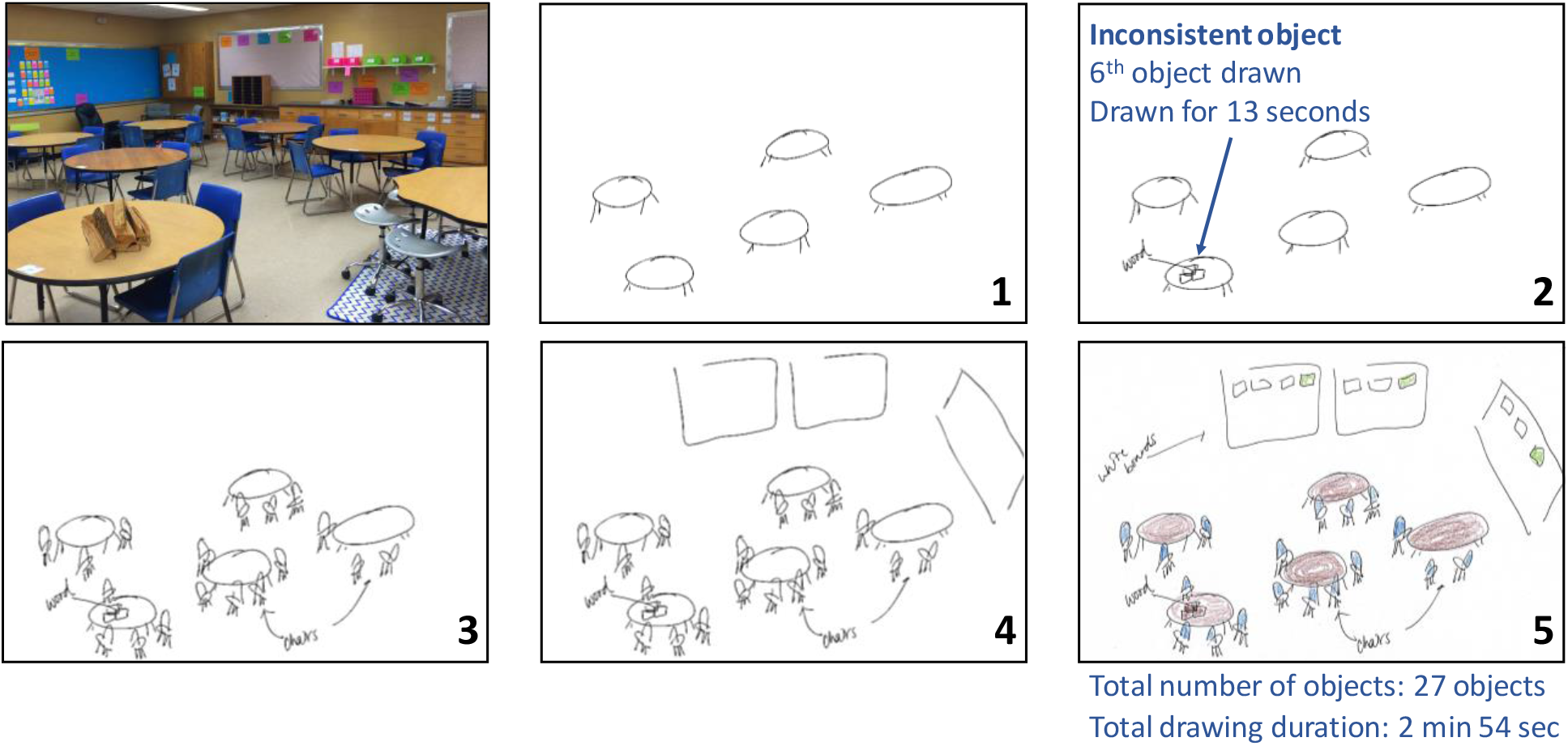
Example of pen tracking output and recall order analyses. For each drawing, the pen tablet recorded a video of the pen strokes in order. This figure shows 5 ordered example frames from one drawing video of an inconsistent classroom scene (with a pile of logs). For the manipulated object, a scorer noted the order in which the object was drawn (normalized by total number of objects), and the length of time it was drawn for (normalized by total time spent on the drawing).

### Post-drawing recognition task performance

Finally, we also tested participants’ memory for the items using a visual recognition task following the drawing recall task. When tested for their recognition of each background scene (with the manipulated object concealed by a gray circle), participants had very high recognition accuracy regardless of whether the scene was originally consistent or inconsistent (Inconsistent: Mean hit rate=92.2%, SD=11.4%; Consistent: M=88.9%, SD=13.4%), with no significant difference between the groups (*t*(29)=1.00, *p*=0.326). There was also no significant difference in false recognitions of matched foil images from the same scene category (Inconsistent: Mean false alarm rate=11.2%, SD=14.8%; Consistent: M=10.0%, SD=11.2%; *t*(29)=0.45, *p*=0.656). Participants then were presented with four possible objects to fill in the obscured part of the image: 1) the inconsistent exemplar, 2) the consistent exemplar, 3) a different exemplar image from the inconsistent object category, 4) a different exemplar image from the consistent object category. There was no significant difference between inconsistent and consistent scenes in participants being able to choose the correct item out of the four options (Inconsistent: Mean hit rate=28.2%, SD=20.9%; Consistent: M=26.5%, SD=14.6%; *t*(28)=0.39, *p*=0.697). There were also no significant differences in the types of errors made by participants. Participants across groups were equally likely to choose an object that appeared in another image (Inconsistent: M=27.1%, SD=19.1%; Consistent: M=23.5%, SD=18.5%; *t*(28)=0.79, *p*=0.434), or an object of the correct category but an incorrect exemplar (Inconsistent: M=18.3%, SD=16.6%; Consistent: M=24.8%, SD=18.3%; *t*(28)=1.34, *p*=0.192).Thus, while we measured differences in recall performance in the drawing task, clear differences in recognition performance did not appear. That being said, participants reported this object recognition task was difficult (occurring after the relatively effortful drawing recall task and with very closely-matched foil objects), and performance was relatively low.

## Discussion

In this study, we tested how memory representations may differ based on consistent or inconsistent object-scene semantics using a visual recall drawing task. We found that scenes containing inconsistent objects were recalled more often, but with less detail. Further, object-scene inconsistencies resulted in a weaker binding between the object and its scene, with the inconsistent object sometimes drawn in isolation, with an unspecified object identity, or transposed into an entirely different scene. In contrast, while semantically consistent scenes were recalled less frequently, their successful recollections contained more object details, and fewer errors.

These results help to reconcile a debate in the field on the impact of object-scene semantics on memory (Friedman, 1979; Pedzek et al., 1979; Hollingworth et al., 2001; Draschkow & Võ, 2017; Cornelissen & Võ, 2017). Using drawing as a memory output allows for a fine-grained look at how object-scene semantics influence memory representations, and we observe a nuanced trade-off in which memory for the overall image is better, but memory for the objects within it is worse. This dual result could account for the fact that some work had previously observed diminished memory for inconsistent images (Draschkow and Võ, 2017) while others observed improved memory (Friedman, 1979; Pedzek, Whetstone, Reynolds, Askari, & Dougherty, 1989; Hollingworth, Williams & Henderson, 2001). In fact, both effects may be occurring simultaneously at different levels of stimulus information (i.e., the image, the objects, and the background scene). This could reflect an effect akin to the trade-off of capacity and precision observed in visual working memory (Roggeman, Klingberg, Feenstra, Compte, & Almeida, 2014), in which inconsistent scene semantics may result in higher capacity with less precision. These results could also provide evidence for the contextual guidance model suggesting two parallel pathways for scene processing: one for gist-based global information and one for object-based local information (Torralba, Oliva, Castelhano, Henderson, 2006; Võ & Wolfe, 2015). While scene-object inconsistencies may result in a distinctive scene with boosted memory for the gist of the scene, they may prevent the ability to use a scene template to fill in local object details (Malcolm & Henderson, 2009; Hollingworth, 2009). A disruption of the scene-object semantics may also result in looser binding of the objects to its scene, resulting in a “spotlighting” on the inconsistent object (Cornelissen & Võ, 2017), and a tendency to migrate objects across memory episodes (Hannigan & Reinitz, 2003). Within memory, semantically inconsistent objects may impair abstraction of the scene from a schema template (Hock & Schmelkopf, 1980; Intraub, 1997), resulting in a loss of schema-coherent details. While the current study focused on the consistency of a single object with its greater scene, investigations of recall for more complex semantic manipulations (e.g., manipulating the semantic relationships of the objects to each other) may provide further insight on how semantics during perception influence the memory representation for a scene.

While we observed differences in recall for consistent and inconsistent scenes, we also observed several similarities between memory representations for consistent and inconsistent scenes. Between these two conditions, recalled drawings tended to be equally diagnostic, have equally high spatial accuracy (in terms of both object location and size), and equally rare numbers of additional objects inserted into the drawings. We also did not observe differences between the two conditions in visual recognition performance (although this could be due to the difficulty of the recognition task). Thus, while scene semantics may influence some aspects of a memory (e.g., memory for other objects in an image), it may have less of a sway on other aspects of that memory (e.g., spatial accuracy). Indeed, various work has suggested differences in how object and spatial information may be coded in memory (Farah & Hammond, 1988; Staresina, Duncan, & Davachi, 2011; Bainbridge, Pounder, Eardley, & Baker, 2019). While the current work investigates scene semantics, other work has suggested that scene *syntactics*–the spatial arrangement of semantically consistent objects within a scene–as similarly meaningful organizational principles for scenes (Võ et al., 2019). An experiment manipulating scene syntactics rather than semantics (e.g., moving a consistent object to an inconsistent location) may result in higher spatial error but preserved object accuracy in memory.

Finally, our results also suggest attention- and fixation-based models may be insufficient models for recall. Here, we are successfully able to replicate findings suggesting that inconsistent objects capture eye movements during perception (Loftus & Mackworth, 1978; De Graef et al., 1990; Henderson et al., 1999; Malcolm & Henderson, 2010). We are also able to replicate prior work showing that computer-vision-based visual saliency metrics can successfully model eye movements on an image (Harel et al., 2007; Kümmerer et al., 2016). While these models do also integrate high-level visual information such as object category (specifically DeepGaze II), it is possible that this ability for computer vision models to predict fixation patterns on the images may indicate there is also a visual difference between inconsistent and consistent images. While we had counterbalanced consistent-inconsistent pairs across participants, it is possible the inconsistent images may have been more visually striking (e.g., a colorful beach ball in a monochromatic laboratory) and captured fixations. However, importantly, we do not find a relationship between fixation patterns during study and pen movement patterns during recall. Further, we do not find a relationship between fixation time or saliency of an object and its probability of being recalled. Thus, our findings are likely due to semantically driven differences in memory rather than visually driven differences. Prior work has found key differences between saliency-based predictions and recall, such as a lower visual field bias for object recall not present in saliency models (Bainbridge et al., 2019) as well as an inability for saliency models to capture semantically meaningful portions of an image (Bylinskii et al., 2016; Henderson & Hayes, 2017). The current work highlights a need for image-based metrics aimed at making predictions specific to scene memory, accounting for semantic abstraction of the scene as well as what objects and features are memorable (Bainbridge, 2019).

In sum, this study reveals a multiple-pronged impact of scene semantics on visual memory representations. These findings are markedly different from previous results showing attentional capture by semantically inconsistent objects, as fixation- and image-based metrics do not show significant relationships with what participants recalled in their drawings. While semantic inconsistencies result in highly atypical images that are remembered overall, these inconsistencies disrupt memory for local object detail in the scenes.

## Acknowledgements

We thank Anna Corriveau for her help digitizing the drawings from the study, and Adam Dickter for his help with the eye tracker system. This research was supported by the Intramural Research Program of the National Institutes of Health (ZIA-MH-002909), under National Institute of Mental Health Clinical Study Protocol 93-M-1070 (NCT00001360). All drawings and online scoring data will be made publicly available on the Open Science Framework (link TBD).

